# A systems-level analysis of dynamic total-body PET data reveals complex skeletal energy metabolism networks *in vivo*

**DOI:** 10.1101/2021.02.16.431368

**Authors:** Karla J. Suchacki, Carlos J. Alcaide-Corral, Samah Nimale, Mark G. Macaskill, Roland H. Stimson, Colin Farquharson, Tom C. Freeman, Adriana A. S. Tavares

## Abstract

Bone is now regarded to be a key regulator of a number of metabolic processes, in addition to the regulation of mineral metabolism. However, our understanding of complex bone metabolic interactions at a systems level remains rudimentary, limiting our ability to assess systemic mechanisms underlying diseases and develop novel therapeutics. *In vitro* molecular biology and bioinformatics approaches have frequently been used to understand the mechanistic changes underlying disease at the cell level, however, these approaches lack the capability to interrogate dynamic multi-bone metabolic interactions *in vivo*. Here we present a novel and integrative approach to understand complex bone metabolic interactions *in vivo* using total-body positron emission tomography (PET) network analysis of murine ^18^F-FDG scans, as a biomarker of glucose metabolism signature in bones. In this report we show that different bones within the skeleton have a unique glucose metabolism and form a complex metabolic network. These data could have important therapeutic implications in the management of the metabolic syndrome and skeletal disease. The application of our approach to clinical and preclinical total-body PET studies promises to reveal further physiological and pathological tissue interactions, which simplistic PET standard uptake values analysis fail to interrogate, extending beyond skeletal metabolism, due to the diversity of PET radiotracers available and under development as well as the advent of clinical total-body PET systems.

**One Sentence Summary:** Bones form a complex metabolic network.

## Main text

To date, advances in molecular imaging, the availability of large human omic datasets and bioinformatic tools have equipped researchers to understand molecular processes that cause disease, and identify and develop new therapeutics *(1)*. However, to understand complex physiological and pathological interactions we need to employ system level approaches. Positron emission tomography (PET) imaging allows for the non-invasive investigation of signalling pathways owing to the radiotracer principle and total-body dynamic PET lends itself to deciphering complex biological processes and interactions *(2, 3)*, such as those found associated with the skeletal system. Most recently, the world’s first clinical total-body PET/Computed Tomography (CT) scanners (PennPET and uExplorer) have been constructed and the first clinical results reveal the potential for rapid scanning, low-dose and whole-body dynamic imaging, which, unlike multi-bed position whole-body PET, allows kinetic analysis in multiple tissues and time points (*4*). Here we present an integrative approach to understand complex tissue interactions *in vivo* using total-body PET network analysis that is directly applicable to emerging clinical total-body dynamic imaging. We initially focused on the skeletal system as it provides an ideal model for analysing complex interactions. The skeleton serves multiple functions *in vivo* such as organ protection, allowing for weight-bearing motion, providing a niche for haematopoiesis and has recently emerged to have major endocrine functions, for example by the bone-specific protein, osteocalcin (*5-8*). Specifically, healthy murine and human bone and cartilage is a significant site of glucose uptake (*7, 9, 10*). However, it remains unclear if different bones within the skeleton have specialised roles in glucose metabolism. Here, we used ^18^F-fluorodeoxyglucose (^18^F-FDG) dynamic total-body PET to explore whether glucose uptake in different bones are associated with one another *in vivo*. Our studies demonstrate that in mice, different bones within the skeleton (e.g. spine) have unique glucose uptake values and form a distinct functional network. Together, these findings underscore the potential for the skeleton to influence metabolic homoeostasis and highlight the importance of studying whole-body physiology using an integrated systems approach.

### Site-specific metabolic differences in bones identified by whole-body PET/CT analysis

The bones of the skeleton are commonly divided into two anatomical classifications; the axial skeleton (bones along the body’s long axis) and the appendicular skeleton (appendages of the axial skeleton). In addition to bone location, bones can form via two fundamentally different processes. Flat bones (e.g. the skull and scapula) are formed by intramembranous ossification, whereas long bones (e.g. tibia and humerus) are formed by endochondral ossification (*11*). Traditionally, dual energy x-ray absorptiometry (DEXA) or quantitative computed tomography (qCT), have been used for quantification of bone density in the clinic (*12, 13*). Although DEXA and qCT data have been used for studying bones’ density in the preclinical and clinical setting for decades, this approach provides a limited window into skeletal function. This needs to be widened owing to recent findings showing the skeleton to be a significant site of glucose uptake and to be involved in the endocrine regulation of whole-body glucose metabolism (*7, 14-16*). To test if individual bones have distinct roles in glucose metabolism we analysed ^18^F-FDG uptake in mice (**Figure 1a, Supplemental Figure 1**) and extracted the standardised uptake values (SUV) from the appendicular (tibia, femur, humerus, ulna and radius (forearm)) and the axial (spine, sternum and skull) skeleton (**Figure 1b-e**). Our PET results in mice showed that overall ^18^F-FDG uptake in the skeleton was bone specific and un-related to bone density measured by qCT. Overall, measured SUVs in the axial skeleton were higher than in the appendicular skeleton while measured Hounsfield Units (HU) from qCT showed higher mineral density in the appendicular skeleton than in the axial skeleton. These findings were congruent throughout the analysis at individual subject-level (heat maps, **Figure 1b**), group averages statistical analysis (Box-and-whisker plots, **Figure 1c-d**) and relative fraction analysis (pie-charts, **Figure 1e**). The qCT data had lower measurement variability and thus outputted several statistically significant differences at group-level analysis (**Figure 1c**). Unsurprisingly, PET data had higher measurement variability owing to the functional nature of the technique, therefore reporting a smaller range of significant differences. Arguably the expected high ^18^F-FDG uptake by the myocardium and brain may have affected SUV quantification in bones in close proximity to these organs (namely sternum and skull) due to spill-over measurement errors. However, these are expected to contribute to a maximum of 10% quantitative bias, as per measurements in our scanner using phantoms and standard acquisition protocols developed by our group (*17*).

**Figure 1.**
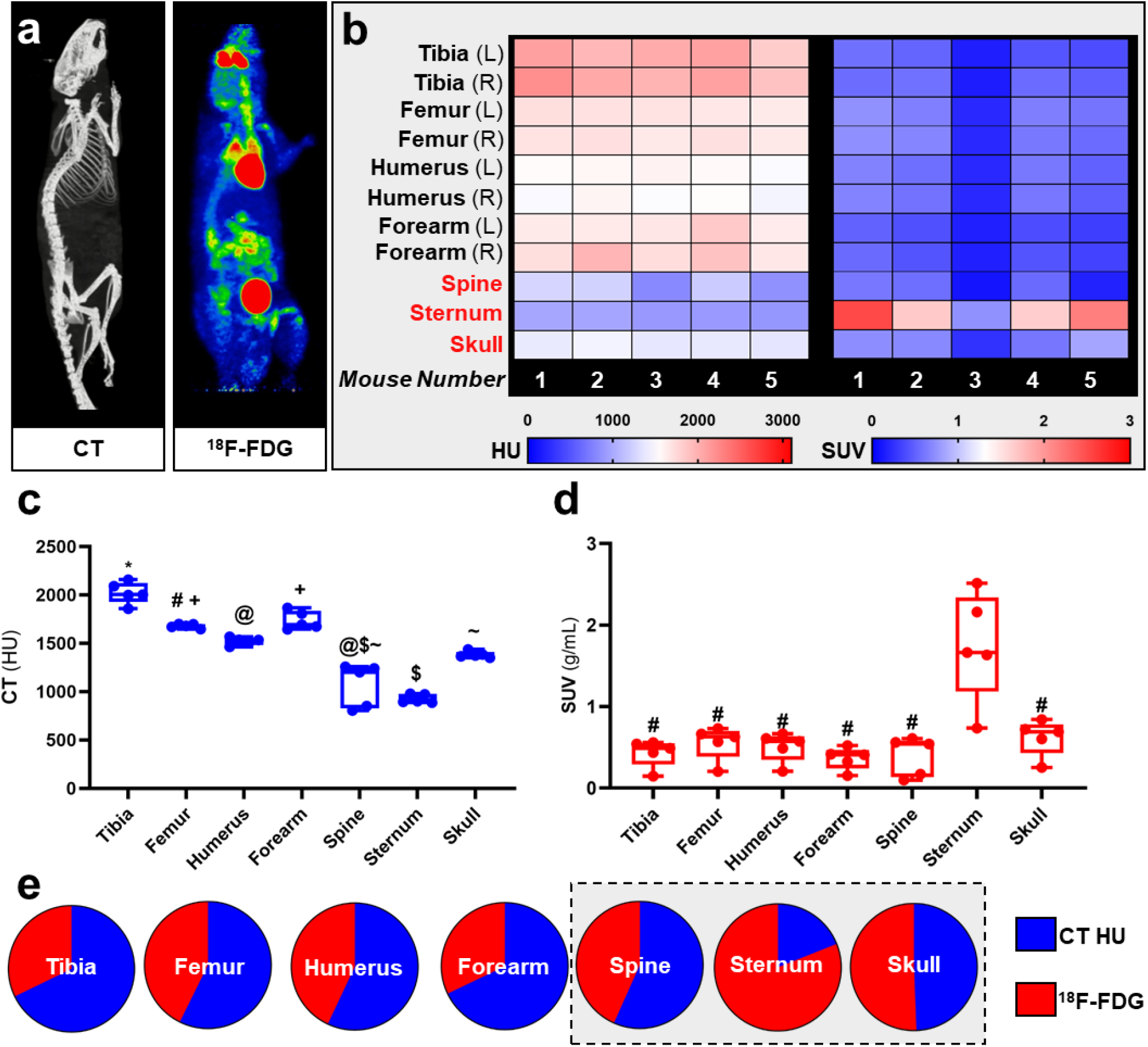
Site-specific metabolic differences in bones identified by whole-body PET/CT analysis. **(a)** Representative maximum intensity projection images of CT and PET data following intravenous administration of ^18^F-FDG. **(b-d)** Hounsfield units (HU) and standard uptake values (SUV) of ^18^F-FDG. SUV was calculated by averaging mouse dynamic PET time-activity curves between 45-60 min post-injection into a single static data point for each bone. **(e)** SUV uptake and HU per bone calculated as a percentage of total ^18^F-FDG and HU, respectively, with axial VOIs (sternum, spine and skull) highlighted in grey. PET SUV percentages were calculated relative to all SUV’s measured in the different bones and then plotted with CT HU percentages calculated relative to all CT HU measured in different bones. Heatmaps are measured SUV and HU of 5 individual mice. Appendicular VOIs (tibia, femur, humerus and forearm) are highlighted by black text and axial VOIs (sternum, spine and skull) are highlighted in red text. Box-and-whisker plots; boxes indicate the 25^th^ and 75^th^ percentiles; whiskers display the range; and horizontal lines in each box represent the median. Significant differences were determined by a one-way ANOVA with multiple comparisons. Different symbols above the error bar show significant difference at *P <* 0.05 **(c)**. # indicates different from sternum at P *<* 0.05 **(d)**.

### Complex bone metabolic network identified by innovative dynamic total-body PET analysis

Having identified site-specific glucose uptake value differences in bones using murine ^18^F-FDG PET data where axial bones have on average higher glucose metabolism than appendicular bones (**Figure 1**), we tested if individual bones’ distinct metabolism formed functional interconnected networks at a system level. A network clustering analysis was performed on the extracted time-activity curves of ^18^F-FDG obtained using dynamic 1-hour total-body PET scanning (**Figure 2a**) to investigate interactions between individual bones and identify if glucose skeletal metabolism networks were present. We identified a unique functional network (**Figure 2b-c**) whereby there was a high connectivity between long bones (femur, tibia). Meanwhile, the spine showed very little connectivity to any other bony tissue in the glucose metabolism (^18^F-FDG) network. Furthermore, the spine, which had the weakest connectivity in the^18^F-FDG skeletal network, was the only bone to show a strong positive correlation (R^2^ = 0.9965) between ^18^F-FDG uptake and bone density (by CT HU, **Figure 2d**). Osteoblasts have been shown to be extremely avid glucose seekers (*18*) and express the glucose transporters Slc2a1, Slc2a4, and Slc2a3 (*19*). These data suggest that the osteoblasts of the spine utilise increased glucose compared to osteoblasts situated at other skeletal sites. However, 90-95% of the cells that reside within the bone matrix are osteocytes (*20*). Derived from terminally differentiated osteoblasts and entombed in the bone matrix, osteocytes sense mechanical stress and communicate signals to initiate or stop the remodelling sequence (*21*). It has recently been shown that glucose levels directly regulate osteocyte function through sclerostin expression (*22*) however little more is known about osteocyte energy status and bioenergetics and it is possible that osteocytes of the spine also utilise increased glucose compared to osteocytes situated at other skeletal sites.

**Figure 2.**
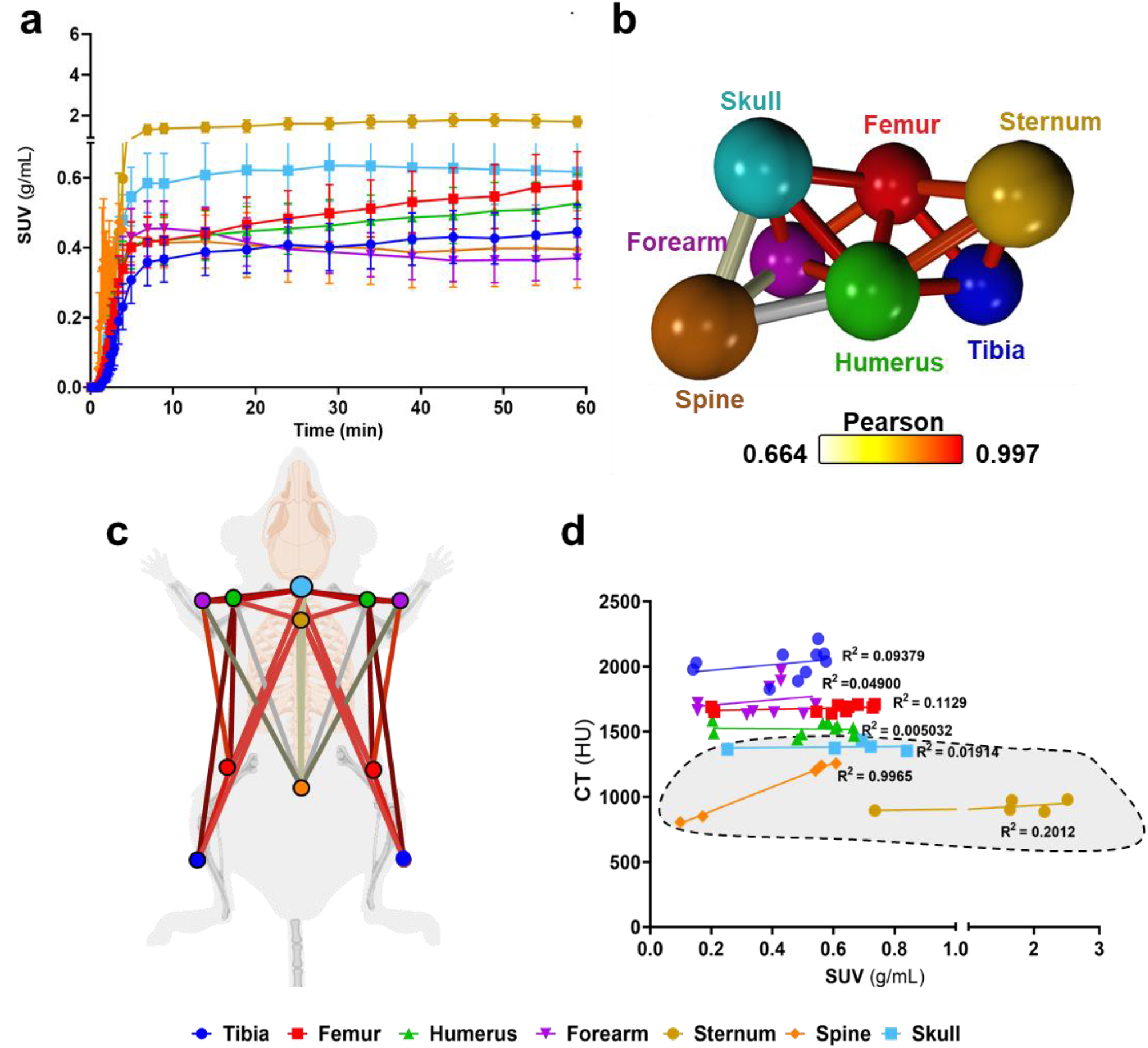
Complex bone metabolic networks identified by innovative dynamic total-body PET network analysis. **(a)** Time-activity curves expressed as standard uptake values (SUV) ^18^F-FDG (n=5). **(b)** Functional network analysis of ^18^F-FDG time-activity curves whereby nodes represent the individual skeletal bones and the edges demote the Pearson correlation value between the nodes (k-nearest neighbors, kNN, of 3). **(c)** Pictorial representation of skeletal networks identified using the functional network analysis of time-activity curves of ^18^F-FDG, nodes are colour coded to represent each bone and the conneting lines demote the Pearson correlation value between the nodes. **(d)** Correlation between computed tomography (CT) Hounsfield units (HU) and SUV from ^18^F-FDG average between 45-60 minutes. Data are presented as the mean ± SEM, n=5. Simple linear regression R^2^ values for VOIs are denoted on d and axial VOIs (sternum, spine and skull) are highlighted in grey.

### Significance of findings and concluding remarks

Our study demonstrates that in mice, different bones within the skeleton have unique molecular signatures and form a distinct metabolic network. Of importance, the metabolic dissimilarity observed between the spine and the rest of the skeleton identified only by the ^18^F-FDG total-body network and not standard whole-body SUV analysis may be of significant clinical importance and could impact on the development of new treatments for metabolic and bone diseases.

Whole-body PET analysis provides one snap-shot of the glucose uptake across multiple tissues at a single time-point. Conversely, total-body PET analysis allows for unprecedented exploration of glycolytic rate kinetics at multiple tissues and time points and how they network to maintain tissue homeostasis at a systems level. Application of the network principles for analysis of PET data, like we presented here, provides a rapid and scalable platform to explore and identify new tissue functions and dysfunctions at system-level that would not be feasible using standard whole-body PET approaches. For example, osteoporosis, a systemic skeletal disease characterised by low bone mass and microarchitectural deterioration is estimated to be responsible for 80–95% of hip and spine fractures in humans *(23, 24)* and is a result of oestrogen deficiency and of the ageing process. In murine models of aging, ^18^F-FDG PET/CT analysis has shown that the spine had reduced ^18^F-FDG uptake compared to other skeletal sites and this uptake was reduced with increasing age *(10).* Using total-body PET/CT data and network analysis, we were able to show the spine was functionally distinct from other bones in the mouse and its density is strongly dependent on glucose metabolism, thus suggesting spine fragility during the aging process and osteoporosis might be underpinned by a stronger dependence on glucose metabolism. Consequently, based on our data, new treatments for bone diseases such as osteoporosis should be carefully designed to account for these skeletal site-specific metabolic differences while keeping in mind systems level interactions beyond bone mineralisation.

Furthermore, the consequences of bone and cartilage disease on energy metabolism regulation remain poorly understood. Importantly, cardiovascular disease is a major cause of premature death and chronic disability (*25*) and musculoskeletal conditions are the second largest contributor to disability worldwide, prevalent in one third to one half of multi-morbidity presentations (*26*). Therefore, the results presented in this study are an important step to unravel these complex systems interactions which standard PET SUV analysis fails to interrogate. Moreover, these data are directly relevant to human health due to the recent development of the first clinical total-body PET systems, which will provide an opportunity to investigate if our findings in mice translate to humans. One can easily envision the application of the innovative total-body PET network analysis technique reported in this paper in a variety of diseases and the characterisation of network changes or losses during pathology, for example, were there is metabolic disruption at system-levels.

In conclusion, we have shown that simplistic CT HU and PET SUV analysis fail to interrogate functional system-level networks that are present *in vivo*. Our novel network-based analyses of PET data have highlighted that the spine has a unique glucose metabolic function where bone density is strongly dependent on glucose metabolism. Applying our new PET network analysis approach to other preclinical studies and clinical studies holds great promise in not only revealing further physiological and pathological intricacies of the skeleton, but can also be used to understand physiological and pathological tissue interactions between organ systems.

## Acknowledgements

AAST is funded by the British Heart Foundation (FS/19/34/34354) and is a recipient of a Wellcome Trust Technology Development Award (221295/Z/20/Z). KJS and RHS are funded by the Medical Research Council (MR/S035761/1) and RHS is funded by the Chief Scientist Office (SCAF/17/02). MGM is funded by the British Heart Foundation (RG/16/10/32375). CF is supported by the Biotechnology and Biological Sciences Research Council (BBSRC) through an Institute Strategic Programme Grant Funding (BB/J004316/1).The British Heart Foundation is greatly acknowledged for providing funding towards establishment of the preclinical PET/CT laboratory (RE/13/3/30183). We thank Mr. William Mungal for invaluable technical assistance with the animal experiments; CJAC is supported by the Edinburgh Preclinical Imaging core facility. We are grateful to Dr Tashfeen Walton and Dr Christophe Lucatelli (Edinburgh Imaging, University of Edinburgh) for radiotracer production. Authors have no competing interests.

## Supplemental materials

### List of Supplementary Materials

Materials and Methods, Supplementary Figure 1.

## Materials and Methods

### Animals and study design

Studies were done in compliance with all relevant ethical regulations under project licences granted by the UK Home Office, and were approved by the University of Edinburgh Animal Welfare and Ethical Review Board. Male 13-week-old C57BL/6JCrl (n=5) mice were bred in-house and housed at 22–23 °C on a 12 h light/dark cycle with free access to water and food, as indicated. The experimental design is outlined in Supplementary Figure 1.

**Supplementary Figure 1.**
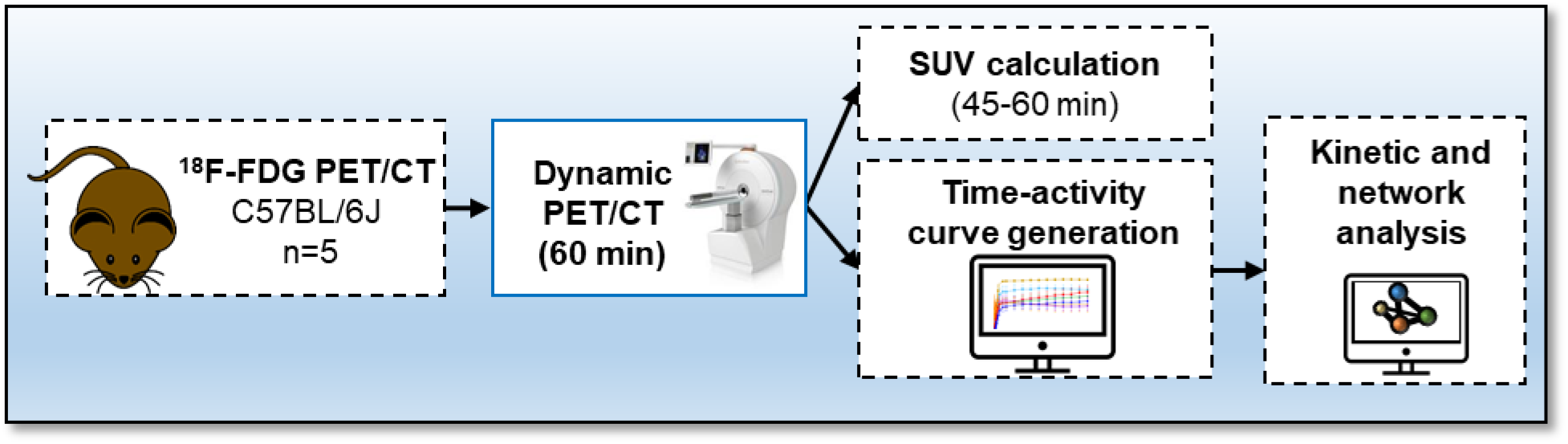
Protocol for ^18^F-fluorodeoxyglucose (FDG) PET/CT. Mice received an intravenous bolus injection via tail-vein of ^18^F-FDG and immediately underwent a 60-min total-body emission scan. A CT scan was conducted at the end of each PET scan. Time activity-curves and standard uptake values were calculated and network analysis was performed to visualise interactions between bones using the Pearson correlation values.

### Imaging data acquisition and reconstruction

Prior to PET/CT imaging, mice were weighed, anesthetised with a pre-set with a mixture of 0.5/0.5 L/min of oxygen/nitrous oxide and 3% isoflurane and transferred to the microPET/CT scanner (nanoPET/CT, Mediso, Hungary). General anaesthesia was maintained throughout the duration of the PET/CT study (0.5/0.5 L/min of oxygen/nitrous oxide and 2 % isoflurane), and vital signs, including temperature and respiration rate, were monitored during the experiments. Animals received a tail vein intravenous bolus injection of ^18^F-fluorodeoxyglucose (^18^F-FDG, 15.08±5.87 MBq, mean±SD; Group 2). ^18^F-FDG was produced in-house (Edinburgh Imaging) using standard methods of radiolabelling (*27*).

Immediately following radiotracer administration, animals underwent a total-body emission scan followed by a CT scan (semi-circular full trajectory, maximum field of view, 360 projections, 50kVp, 300ms and 1:4 binning). Collected PET images underwent attenuation correction using the CT data. PET images were reconstructed between 0-60 min into 6×30sec, 3×60sec, 2×120sec and 10×300sec frames using Mediso’s iterative Tetra-tomo 3D reconstruction algorithm and the following settings: 4 iterations, 6 subsets, full detector model, low regularisation, spike filter on, voxel size 0.4 mm and 400-600 keV energy window. PET data were corrected for random coincidences, scatter and attenuation.

### Image processing and standard uptake value (SUV) calculation

Reconstructed images were analysed using PMOD 3.7 software (PMOD Technologies, Switzerland). Volumes of interest (VOI) were drawn around the tibiae, femurs, humeria, radius and ulnas (forearm), spine, sternum and skull. To distinguish bone tissue from bone marrow and surrounding tissues, the VOIs were segmented using previously defined Hounsfield Units, HU, (332-50000) generated using HU obtained from the acquisition of a CT tissue equivalent material (TEM) phantom (CIRS, model 091) and mouse CT scans (*7*). All dynamic PET data were then corrected for time delays between start of the scan and injection of radiotracer. Time activity-curves were generated and standard uptake values (SUVs) were calculated by normalising radioactive concentration in VOI for the injected dose and the animal weight. To estimate the bone uptake at equilibrium, SUV averages were taken from 3 PET frames between 45-60 min post-radiotracer administration. The CT HU were extracted from the VOI of the tibiae, femurs, humeria, forearm spine, sternum and skull.

### Network analysis of total-body PET data

Non-decay corrected dynamic total-body PET data was summarised into a table with rows representing radiotracer signal from an individual bones per study group (1 and 2) and columns the time of the recording. The file was saved as a comma separated variable (.csv) file. This was loaded in the network analysis tool Graphia (https://graphia.app/) (*28*). Pearson correlations were calculated for Group 1 and 2 and a relationship matrix graph was constructed by performing an all versus all comparison of the ^18^F-FDG signal profiles from each bone (correlation cut off value of R>0.7). By minimising the number of edges the structure of the relationship between tissue-accumulation profiles are revealed, as reflected by graph’s structure and edge weights, where the nodes represent each bone and edges represent correlations above the selected threshold, where the threshold value was set to maintain the number of nodes in the network hence all available data.

### Statistical analysis, data presentation and reproducibility

^18^F-FDG SUV averages were analysed for normal distribution using the Shapiro-Wilk normality test. Simple multiple linear regression was conducted to assess CT and SUV correlations. Data are represented as the average ± SEM, unless otherwise stated in the results section. All statistical analysis was performed using Prism 8 (GraphPad v8, USA). Mouse cartoon networks were created with BioRender.com

## Notes

### Competing Interest Statement

The authors have declared no competing interest.

